# Sexual dimorphism of synaptonemal complex length among populations of threespine stickleback fish

**DOI:** 10.1101/2020.07.30.228825

**Authors:** Barquiesha S. Madison, Shivangi Nath, Mary K. Flanagan, Brittany A. Dorsey, Michael A. White

## Abstract

Crossover frequencies often differ substantially between sexes (i.e. heterochiasmy). Although this phenomenon is widespread throughout taxa, the mechanisms that lead to heterochiasmy remain unclear. One pattern that has emerged is that the overall length of the synaptonemal complex likely has a direct influence on the total number of crossovers in each sex. However, this has only been investigated in a handful of species. The threespine stickleback fish (*Gasterosteus aculeatus*) is an excellent species to explore whether synaptonemal complex length is associated with differences in the total number of crossovers, as females have much longer linkage maps than males. We used an immunocytological approach to quantify synaptonemal complex length in late pachytene female and male meiocytes in two different populations of threespine stickleback fish. Overall, the freshwater population had shorter synaptonemal complex lengths than the marine population. In both populations we observed sexual dimorphism, with females possessing longer axes. Our results support a model where chromosome axis length determines overall crossover frequency and establish the threespine stickleback as a useful species to explore the mechanistic basis of heterochiasmy as well as the genetic basis underlying variation in synaptonemal complex length.

## Introduction

Crossing over is an essential step of meiosis that is required for the proper pairing and segregation of homologous chromosomes. At minimum, a single crossover is typically required per chromosome or chromosome arm for meiosis to complete correctly (Villena and Sapienza 2001; Sun *et al*. 2004; Segura *et al*. 2013; Dumont 2017). Errors during crossover formation may lead to severe segregation errors resulting in aneuploidies (Hassold and Hunt 2001). Beyond this minimum requirement, the overall frequency at which this essential process occurs throughout the genome can vary tremendously. Genome-wide rates of crossing over are highly variable between species and even within species (reviewed in Coop and Przeworski 2007; Smukowski and Noor 2011; Ritz *et al*. 2017), with some of the greatest differences occurring between sexes (i.e. heterochiasmy; reviewed in Sardell and Kirkpatrick 2019).

The mechanism that leads to rate differences between sexes is unclear; however, the chromatin environment imposed by the synaptonemal complex (SC) likely plays a role. The SC is a proteinaceous axis that links homologous chromosomes together at mid-prophase (reviewed in Page and Hawley 2004). This structure serves as a scaffold in which DNA double strand breaks are repaired between homologous chromosomes in a crossover or non-crossover fashion. There is a strong positive correlation between crossover number and the total length of the SC (Lynn *et al*. 2002; Tease and Hultén 2004; Drouaud *et al*. 2007; Kochakpour and Moens 2008; Giraut *et al*. 2011; Gruhn *et al*. 2013; Fröhlich *et al*. 2015; Ruiz-Herrera *et al*. 2017), suggesting that this structure may directly determine the overall frequency of crossover events. Even among individuals within a species, SC length can vary widely (Pan *et al*. 2012; Wang *et al*. 2019), altering the organization of DNA along the chromosome axis (Blat *et al*. 2002; Kleckner *et al*. 2003; Kauppi *et al*. 2012). The same length of DNA organized along a longer chromosome axis, results in smaller chromatin loops and a higher frequency of crossover formation.

Heterochiasmy is correlated with sex-specific differences in SC length of some species. In mammals, females exhibit longer SC complexes, coincident with higher crossover frequencies, relative to males (Lynn *et al*. 2002; Tease and Hultén 2004; Gruhn *et al*. 2013). Heterochiasmy is widespread in plants and animals (reviewed in Sardell and Kirkpatrick 2019), yet it remains unknown whether altered SC length is a universal mechanism responsible for this phenomenon. Outside of mammals, dimorphic SC lengths have only been noted in teleost fish, and these analyses have revealed conflicting patterns. Zebrafish have been shown to have longer SC lengths in female fish, congruent with a higher frequency of crossing over (Kochakpour and Moens 2008). However, in cichlid fish there has been no consistent pattern identified between SC length and the sex-specific frequency of crossing over (Campos-Ramos *et al*. 2009). Additional teleost species are needed to assess the generality of these mechanisms.

Similar to other organisms, threespine stickleback fish (*Gasterosteus aculeatus*) exhibit striking dimorphism in crossover frequency between males and females. Sex-specific linkage maps in this species have shown females have crossover frequencies that are 1.6 times higher relative to male fish (Sardell *et al*. 2018). Despite these differences in crossover frequency, there is no evidence of dimorphic SC lengths. Chromosome axis lengths have been measured in a single freshwater river population of fish using electron microscopy (Cuñado *et al*. 2002). In this population, males and females had axes of similar length at pachytene. Because these two studies used different source populations, it raises the possibility that SC length dimorphism may be polymorphic, increasing female crossover frequency in select populations. Alternatively, there may not be sexual dimorphism of SC length in threespine stickleback fish, and heterochiasmy may be driven by an independent mechanism. Here, we use immunofluorescence to characterize male and female SC length in a freshwater and a marine population of threespine stickleback fish. Unlike the previous study, we document a significantly longer SC in females in both populations, relative to males. Our results align with sex-specific linkage maps in threespine stickleback (Sardell *et al*. 2018) and suggest changes in SC length may be responsible for the altered crossover landscape between sexes in some populations.

## Materials and Methods

### Ethics statement

All procedures using threespine stickleback fish were approved by the University of Georgia Animal Care and Use Committee (protocol A2018 10-003-Y2-A5).

### Preparation of meiotic chromosome spreads from threespine stickleback fish

Meiotic chromosome spreads were isolated from male and female threespine stickleback fish following previously established protocols from mice (Dia *et al*. 2017) and teleost fish (Araya-Jaime *et al*. 2015). Threespine stickleback fish undergo seasonal gametogenesis (Craig-Bennett 1931; Borg 1982; Borg and Veen 1982). Therefore, we targeted stages of juvenile development when meiosis was actively occurring in male and female fish. This was determined to occur between five and six months after hatch. The gonads were dissected from five male fish and five female fish, isolated from a lab marine population of wild-caught fish from Port Gardner Bay (Everett, Washington, USA) and one male fish and four female fish from a lab freshwater population of wild-caught fish from Lake Washington (Seattle, Washington, USA). The gonads were immersed in 500 μl of phosphate-buffered saline (PBS), and minced into a fine cell solution using a dounce homogenizer with a small clearance pestle. To account for reduced meiocyte populations in females, all ovaries were pooled into a single population sample for homogenization. Testes were homogenized one sample per male. The cell solution was centrifuged at 500 xg for five minutes to pellet the cells. After centrifugation, the supernatant was discarded, and the pellet was resuspended in one of two hypotonic solutions. For females in both populations and males of the Lake Washington population, we used 100 μl of 0.8% sodium citrate and the suspension was incubated at room temperature for twenty minutes. For males from the Port Gardner Bay population, we used 100 μl of 100 mM sucrose and the suspension was incubated at room temperature for twenty minutes. The different hypotonic solutions were used because 0.8% sodium citrate did not result in clear chromosome spreads in males for the Port Gardner Bay population. Importantly, we do not detect an effect of the hypotonic solution on synaptonemal complex length (Figure S1). After incubating, 20 μl of the sodium citrate hypotonic solution was placed and mixed on a microscope slide with 20 μl of sucrose. For the sucrose hypotonic solution, only 20 μl of the cell solution was placed on a microscope slide. Slides from both sexes were fixed by adding 100 μl of 1% paraformaldehyde with 0.15% TritonX-100 and were incubated at room temperature for 2.5 hours in a covered, humidified chamber. After incubating, we removed the slides from the humidified chamber and allowed them to dry overnight. After drying, the slides were washed three times with a fresh solution of 1X PBS (15 minutes, 10 minutes, then 5 minutes). The slides were air-dried and immediately processed for immunofluorescence or stored at −20°C. Stored slides were all used within two weeks, the time period where they produce optimal results (Dia *et al*. 2017).

### Immunofluorescence of axial elements of the synaptonemal complex

We visualized the localization of a cohesin protein (structural maintenance of chromosomes 3; SMC3) within the axial elements that associate sister chromatids (reviewed in Bolcun-Filas and Handel 2018). Samples were first permeabilized for 20 minutes using permeabilization solution (1X PBS, 1mM EDTA, and 1% TritonX-100) (Dawe *et al*. 2018). The solution was drained from the slide and 1.5 ml of 1X antibody dilution buffer (1X PBS, 0.25% normal goat serum, 0.003 g/ml BSA, and 0.005% TritonX-100) was added and incubated for 20 minutes (Dumont and Payseur 2011). A 1:100 dilution of ice-cold primary anti-SMC3 (Abcam; ab9263; Reagent Table) was added onto the slide and sealed with a coverslip and rubber cement. The primary antibody was incubated at 4°C overnight. The coverslip was removed by immersion in 1X PBS. 1.5ml of 1X antibody dilution buffer was added to the slide and incubated for 20 minutes. After incubation, a 1:100 dilution of goat anti-rabbit 488-secondary (Abcam; ab150077; Reagent Table) was added, the slide was sealed with a coverslip and rubber cement, and the sealed slide was incubated at 37°C for two to three hours. Slides were washed three times in 1X PBS. After the slides were washed, 40 μl of 1X 4’,6-diamidino-2-phenylindole (DAPI; Biotium; Reagent Table) was added and the slides were sealed with a coverslip and clear nail polish.

### Imaging and measuring synaptonemal complex length

Cells were visualized using a Leica DM6000 B upright microscope at 63x magnification with DAPI and FITC filter sets. Images were captured using a Hamamatsu ORCA-ER digital camera. A total of 41 oocytes and 134 spermatocytes were determined to be within prophase I of meiosis. We staged each meiocyte to their corresponding substage of prophase I according to the overall organization of the synaptonemal complex, revealed by SMC3 localization. Spreads showing sparse SMC3 localization were characterized as leptotene, spreads with fully formed SMC3 axes, but did not show fully synapsed homologs (having greater than a haploid chromosome count of 21) were characterized as zygotene, and spreads with fully formed SMC3 axes and a haploid chromosome count of 21 were characterized as late pachytene. Only male and female cells that were in late pachytene were used to quantify the synaptonemal complex lengths. At this stage, the synaptonemal complex is fully formed and chromosomes are completely paired.

We measured synaptonemal complex length within oocytes and spermatocytes that were staged to late pachytene. The length of each synaptonemal complex was measured in pixels using the freehand line tool and measure function of ImageJ software (Abràmoff *et al*. 2004; Schneider *et al*. 2012). Each chromosome was measured three times and the average length was used. We were blinded to the sex of each sample before measuring. Pixel-based lengths of each chromosome were converted to microns based on a conversion factor specific to the digital camera and camera adapter.

### Data Availability

Molecular cytology images are available upon request. Tables S1 through S4 contain the total SC lengths per meiocyte and the SC lengths per chromosome for each population. The supplemental tables are available through figshare. The authors affirm all other data necessary for confirming the conclusions of this article are present within the article and figures.

## Results and Discussion

### Males and females exhibit fully synapsed chromosomes at pachytene

Threespine stickleback fish have heteromorphic X and Y chromosomes that have diverged from one another (Peichel *et al*. 2004, 2020; Ross and Peichel 2008; White *et al*. 2015). In some species with heteromorphic sex chromosomes, synapsis is limited to the short pseudoautosomal region, leaving the remainder unpaired between homologs (Solari 1970, 1974; Moses *et al*. 1975). We therefore first wanted to explore whether males exhibit any differences in synapsis among chromosomes. We targeted structural maintenance of chromosomes 3 (SMC3), a protein important for the cohesion of sister chromatids during meiosis (reviewed in Bolcun-Filas and Handel 2018) and characterized the different substages of prophase I in a marine population of fish (Port Gardner Bay). In total, we staged 134 spermatocytes and 41 oocytes to leptotene, zygotene, or late pachytene (Figure 1). From the total pool, 16 spermatocytes and 11 oocytes were in late pachytene of meiosis I. We found meiocytes where males and females had a total of 21 fully synapsed chromosomes, consistent with the haploid chromosome count in threespine stickleback fish (Chen and Reisman 1970). Our results indicate the X and Y chromosomes must be fully synapsing in a non-homologous fashion within a large proportion of meiocytes, similar to what was previously observed using electron microscopy (Cuñado *et al*. 2002).

**Figure 1.**
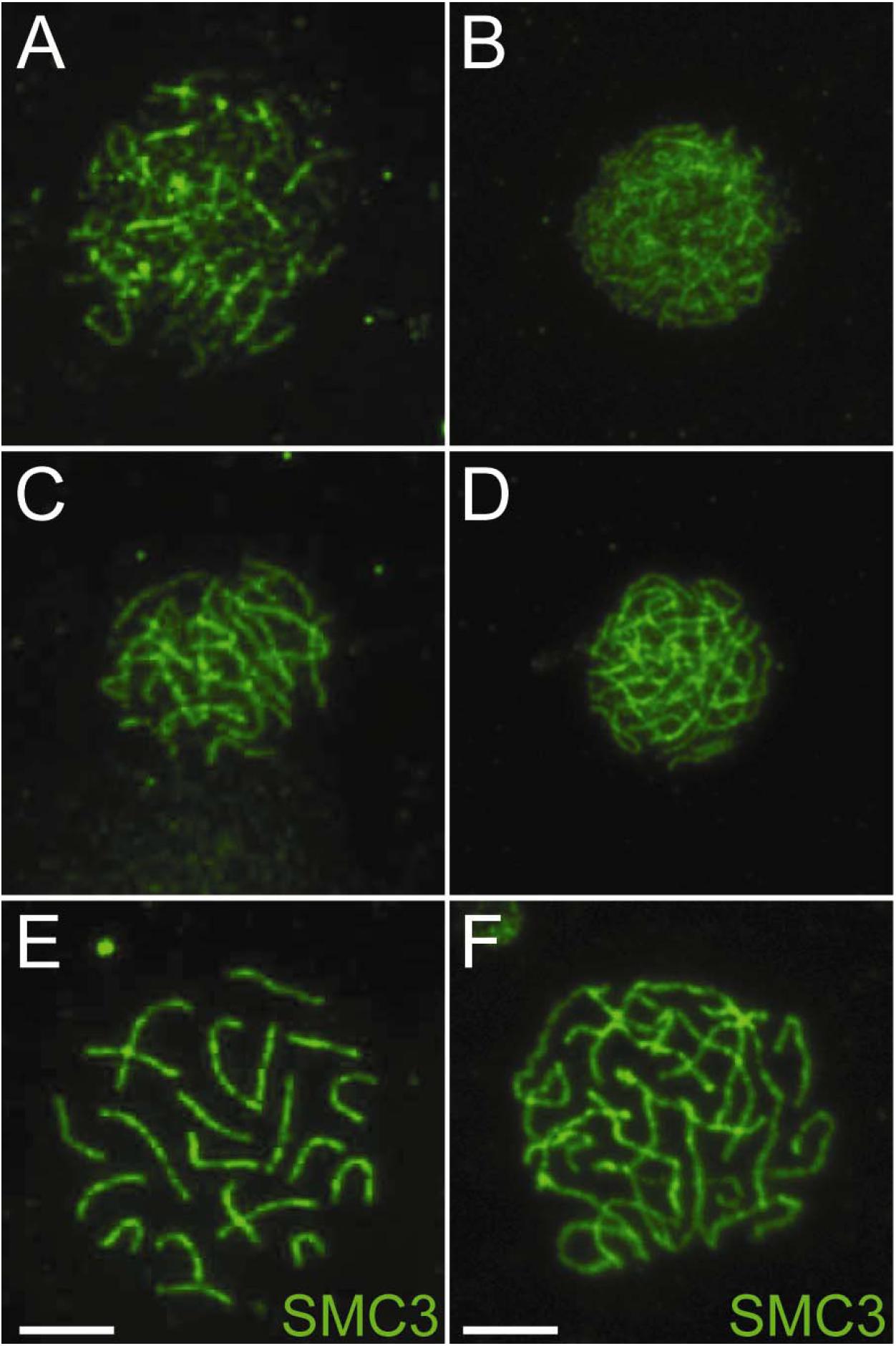
Progression of synapsis throughout prophase in female and male meiocytes. Axial elements of the synaptonemal complex are labeled for localization of the cohesion protein, SMC3 during (A) male leptotene, (B) female leptotene, (C) male zygotene, (D) female zygotene, (E) male pachytene, and (F) female pachytene. Scale bars represent 5 μm.

Highly degenerate, ancient sex chromosomes exhibit delayed pairing and double strand DNA break repair during meiosis due to an overall lack of sequence homology between the X and Y chromosomes (Inagaki *et al*. 2010; Kauppi *et al*. 2011; Checchi and Engebrecht 2011). Recently derived sex chromosomes have higher sequence homology, which could facilitate full synapsis. For example, the young X and Y chromosomes of guppy fish fully synapse, albeit in a delayed fashion relative to autosomes (Lisachov *et al*. 2015). The threespine stickleback Y chromosome also has higher sequence homology between the X and Y chromosomes compared to the ancient sex chromosomes of mammals (White *et al*. 2015; Peichel *et al*. 2020). However, there is also substantial structural divergence between the X and Y, with at least three major inversions that disrupt sequence collinearity between the two chromosomes (Ross and Peichel 2008; Peichel *et al*. 2020). Our results indicate the threespine stickleback X and Y must be synapsing in a non-homologous fashion outside of the pseudoautosomal region.

### Females have longer synaptonemal complexes at pachytene

Female threespine stickleback fish have longer genetic maps, relative to males (Sardell *et al*. 2018). In many species, sex-specific crossover frequencies are highly correlated with overall synaptonemal complex length at pachytene (Lynn *et al*. 2002; Singer *et al*. 2002; Tease and Hultén 2004; Kochakpour and Moens 2008; Gruhn *et al*. 2013). We investigated whether threespine stickleback fish also exhibit sexual dimorphism of SC length by measuring SMC3 labeled chromosome axes in the same marine population of fish as well as a second freshwater population (Lake Washington). In both populations, we found clear dimorphism in SC length, where female fish had significantly longer total axis lengths at pachytene compared to males (Port Gardner Bay: 1.65x longer in females; Lake Washington: 1.32x longer in females; Figure 2; Tables S1 and S2; Mann-Whitney U test, *P* < 0.001 between males and females in both populations). There was a greater variability among meiocytes in females, relative to males (Port Gardner Bay female SD: 25.562, Port Gardner bay male SD: 15.718; Lake Washington female SD: 29.141, Lake Washington male SD: 10.973). The total length difference does not appear to be driven by a small subset of chromosomes, as the increase in synaptonemal complex length was still highly apparent when assayed per chromosome (Figure 2; Tables S3 and S4; t-test, *P* < 0.001).

**Figure 2.**
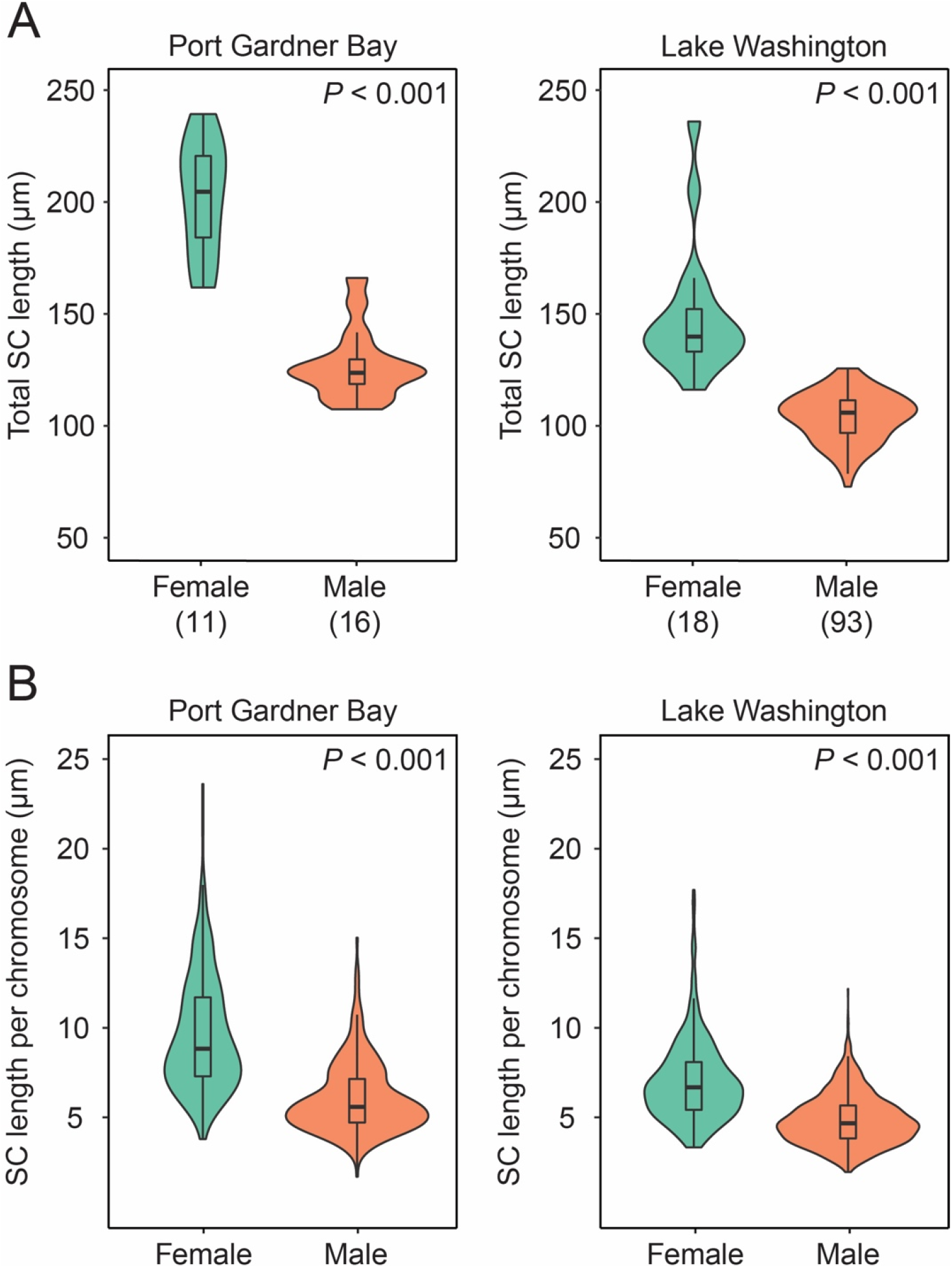
Females have longer synaptonemal complexes than males at late pachytene in two populations of fish. (A) Total length of SMC3 label along the synaptonemal complex was measured at late pachytene. The total number of meiocytes is indicated below. (B) The length of each chromosome is plotted for all meiocytes combined. Whiskers denote 1.5x the interquartile range.

Sex-specific crossover frequency is highly correlated with the overall length of the SC in mammals and zebrafish (Lynn *et al*. 2002; Singer *et al*. 2002; Tease and Hultén 2004; Kochakpour and Moens 2008; Gruhn *et al*. 2013). Our results indicate this correlation also holds in threespine stickleback fish. The SC length difference we observed in both the freshwater and marine population was within range of the difference observed in sex-specific genetic maps in threespine stickleback fish (1.65x longer SC in Port Gardner Bay females; 1.32x longer SC in Lake Washington females; 1.64x longer genetic map in females (Sardell *et al*. 2018)). In addition, we observed population-specific variation in overall SC length. There were significantly shorter SC lengths in both males (Mann-Whitney U test, *P* = 0.006) and females (Mann-Whitney U test, *P* < 0.001) within the freshwater population (Lake Washington), compared to the marine population (Port Gardner Bay). If SC length is correlated with genetic map length, we would expect a corresponding shortening of the genetic map of Lake Washington fish. Consistent with our results, population-scaled crossover rates are significantly lower within Lake Washington compared to another neighboring marine population (Shanfelter *et al*. 2019). Our results highlight SC length is polymorphic among threespine stickleback populations, suggesting crossover rate variation may evolve through changes in this axis. Indeed, variation in SC length has evolved over short time scales between closely related species (Lynn *et al*. 2002; Campos-Ramos *et al*. 2009; Fröhlich *et al*. 2015; Ruiz-Herrera *et al*. 2017; Wang *et al*. 2019)

Our results contrast previous observations in a freshwater river population of threespine stickleback fish where sexual dimorphism was not observed (Cuñado *et al*. 2002). Males in the river population had a mean SC length (150 ± 18 μm) that fell within the range of our measurements. However, mean SC length of females (143 ± 12 <u>μm) fell outside of the range we observed in the marine population. Interestingly, the river population had a female mean that was closer to the length we observed within the Lake Washington freshwater population. This suggests there</u> is polymorphism among populations of threespine stickleback in overall SC length and the genetic architecture controlling this trait may differ between the sexes. In house mice, within species variation in male SC length has been associated with multiple quantitative trait loci (Wang *et al*. 2019). If threespine stickleback fish have a similar genetic architecture shaping SC length, alternate alleles at these loci may have diverged between populations, leading to the observed polymorphism.

Sex-specific crossover frequency can also vary at a fine scale throughout the genome. Males often exhibit a higher frequency of crossovers towards the ends of chromosomes, whereas females have a more uniform distribution (Broman *et al*. 1998; Singer *et al*. 2002; Rexroad *et al*. 2008; Paigen *et al*. 2008; Wong *et al*. 2010; Paigen and Petkov 2010; Giraut *et al*. 2011; Jones *et al*. 2013; Brelsford *et al*. 2016; Smeds *et al*. 2016). An outstanding question is whether these regional shifts in crossover frequency between sexes are also associated with sexual dimorphism of SC length. The only regional difference in SC length between sexes has been examined in the house mouse pseudoautosomal region (Acquaviva *et al*. 2020), the small region of sequence homology between the X and Y that experiences exceptionally high rates of crossing over in males (Soriano *et al*. 1987). The pseudoautosomal region has an extended chromosome axis compared to the ends of some autosomes (Kauppi *et al*. 2011); however, this extended axis has also been found in female meiocytes, where recombination on the X chromosome is not restricted to the pseudoautosomal region (Acquaviva *et al*. 2020). This makes it unclear whether dimorphic SC lengths are responsible for heterochiasmy across specific regions of chromosomes. Threespine stickleback also exhibit heterochiasmy at a fine scale (Sardell *et al*. 2018). They will therefore be a useful species to further explore the mechanisms leading to regional differences in heterochiasmy along chromosomes.

## Conclusions

Our results reveal threespine stickleback fish exhibit differences in SC length among populations of fish and also exhibit sexual dimorphism. This supports a model where sexually dimorphic crossover rates in genetic maps are a result of the chromatin environment imposed by the SC. Future work focusing on variation among populations will shed light on the genetic basis of SC length variation and its relationship with crossover frequency.

## Acknowledgements

The authors thank Lucille Welch for help imaging the female meiocytes. The authors acknowledge the following funding sources: National Science Foundation MCB 1943283 (MAW), Office of the Vice President of Research at the University of Georgia (MAW), and the Rosemary Grant Advanced award from the Society for the Study of Evolution (SN).

## Conflicts of Interest

The authors declare no conflicts of interest.

## Notes

### Competing Interest Statement

The authors have declared no competing interest.

## References

Abràmoff, M. D., P. J. Magalhães, and S. J. Ram, 2004 Image Processing with ImageJ. Biophotonics International 11: 36–42.

Acquaviva, L., M. Boekhout, M. E. Karasu, K. Brick, F. Pratto et al., 2020 Ensuring meiotic DNA break formation in the mouse pseudoautosomal region. Nature 582: 426–431.

Araya-Jaime, C., É. A. Serrano, D. M. Z. de A. Silva, M. Yamashita, T. Iwai et al., 2015 Surface-spreading technique of meiotic cells and immunodetection of synaptonemal complex proteins in teleostean fishes. Molecular Cytogenetics 8: 4.

Blat, Y., R. U. Protacio, N. Hunter, and N. Kleckner, 2002 Physical and functional interactions among basic chromosome organizational features govern early steps of meiotic chiasma formation. Cell 111: 791 802.

Bolcun-Filas, E., and M. A. Handel, 2018 Meiosis: the chromosomal foundation of reproduction. Biol Reprod 99: 112 126.

Borg, B., 1982 Seasonal effects of photoperiod and temperature on spermatogenesis and male secondary sexual characters in the three-spined stickleback, Gasterosteus aculeatus L. Canadian Journal of Zoology 60: 3377 3386.

Borg, B., and T. V. Veen, 1982 Seasonal effects of photoperiod and temperature on the ovary of the three-spined stickleback, Gasterosteus aculeatus L. Can J Zool 60: 3387–3393.

Brelsford, A., N. Rodrigues, and N. Perrin, 2016 High□density linkage maps fail to detect any genetic component to sex determination in a Rana temporaria family. J Evolution Biol 29: 220–225.

Broman, K. W., J. C. Murray, V. C. Sheffield, R. L. White, and J. L. Weber, 1998 Comprehensive Human Genetic Maps: Individual and Sex-Specific Variation in Recombination. Am J Hum Genetics 63: 861–869.

Campos-Ramos, R., S. C. Harvey, and D. J. Penman, 2009 Sex-specific differences in the synaptonemal complex in the genus Oreochromis (Cichlidae). Genetica 135: 325–332.

Checchi, P. M., and J. Engebrecht, 2011 Heteromorphic sex chromosomes: navigating meiosis without a homologous partner. Molecular reproduction and development 78: 623–632.

Chen, T. R., and H. M. Reisman, 1970 A comparative chromosome study of the North American species of sticklebacks (Teleostei: Gasterosteidae). Cytogenetics 9: 321–332.

Coop, G., and M. Przeworski, 2007 An evolutionary view of human recombination. Nat Rev Genet 8: 23–34.

Craig-Bennett, A., 1931 The reproductive cycle of the three-spined stickleback, Gasterosteus aculeatus, Linn. Phil. Trans. Roy. Soc. B. 219: 197 279.

Cuñado, N., J. Barrios, E. S. Miguel, R. Amaro, C. Fernández et al., 2002 Synaptonemal Complex Analysis in Oocytes and Spermatocytes of Threespine Stickleback Gasterosteus Aculeatus (Teleostei, Gasterosteidae). Genetica 114: 53–56.

Dawe, R. K., E. G. Lowry, J. I. Gent, M. C. Stitzer, K. W. Swentowsky et al., 2018 A Kinesin-14 Motor Activates Neocentromeres to Promote Meiotic Drive in Maize. Cell 173: 839–850.e18.

Dia, F., T. Strange, J. Liang, J. Hamilton, and K. M. Berkowitz, 2017 Preparation of Meiotic Chromosome Spreads from Mouse Spermatocytes. J Vis Exp Jove 129: e55378.

Drouaud, J., R. Mercier, L. Chelysheva, A. Bérard, M. Falque et al., 2007 Sex-Specific Crossover Distributions and Variations in Interference Level along Arabidopsis thaliana Chromosome 4. Plos Genet 3: e106.

Dumont, B. L., 2017 Variation and Evolution of the Meiotic Requirement for Crossing Over in Mammals. Genetics 205: 155–168.

Dumont, B. L., and B. A. Payseur, 2011 Genetic analysis of genome-scale recombination rate evolution in house mice. PLoS Genetics 7: e1002116.

Fröhlich, J., M. Vozdova, S. Kubickova, H. Cernohorska, H. Sebestova et al., 2015 Variation of Meiotic Recombination Rates and MLH1 Foci Distribution in Spermatocytes of Cattle, Sheep and Goats. Cytogenet Genome Res 146: 211–221.

Giraut, L., M. Falque, J. Drouaud, L. Pereira, O. C. Martin et al., 2011 Genome-Wide Crossover Distribution in Arabidopsis thaliana Meiosis Reveals Sex-Specific Patterns along Chromosomes. Plos Genet 7: e1002354.

Gruhn, J. R., C. Rubio, K. W. Broman, P. A. Hunt, and T. Hassold, 2013 Cytological Studies of Human Meiosis: Sex-Specific Differences in Recombination Originate at, or Prior to, Establishment of Double-Strand Breaks. PLoS ONE 8: e85075.

Hassold, T., and P. Hunt, 2001 To err (meiotically) is human: the genesis of human aneuploidy. Nature Reviews Genetics 2: 280 291.

Inagaki, A., S. Schoenmakers, and W. M. Baarends, 2010 DNA double strand break repair, chromosome synapsis and transcriptional silencing in meiosis. Epigenetics 5: 255–266.

Jones, D. B., D. R. Jerry, M. S. Khatkar, H. W. Raadsma, and K. R. Zenger, 2013 A high-density SNP genetic linkage map for the silver-lipped pearl oyster, Pinctada maxima: a valuable resource for gene localisation and marker-assisted selection. BMC Genomics 14: 810.

Kauppi, L., M. Barchi, F. Baudat, P. J. Romanienko, S. Keeney et al., 2011 Distinct Properties of the XY Pseudoautosomal Region Crucial for Male Meiosis. Science 331: 916–920.

Kauppi, L., M. Jasin, and S. Keeney, 2012 The tricky path to recombining X and Y chromosomes in meiosis. Annals of the New York Academy of Sciences 1267: 18 23.

Kleckner, N., A. Storlazzi, and D. Zickler, 2003 Coordinate variation in meiotic pachytene SC length and total crossover/chiasma frequency under conditions of constant DNA length. Trends Genet 19: 623–628.

Kochakpour, N., and P. B. Moens, 2008 Sex-specific crossover patterns in Zebrafish *(Danio rerio)*. Heredity 100: 489 495.

Lisachov, A. P., K. S. Zadesenets, N. B. Rubtsov, and P. M. Borodin, 2015 Sex chromosome synapsis and recombination in male guppies. Zebrafish 12: 174 180.

Lynn, A., K. E. Koehler, L. Judis, E. R. Chan, J. P. Cherry et al., 2002 Covariation of Synaptonemal Complex Length and Mammalian Meiotic Exchange Rates. Science 296: 2222 2225.

Moses, M. J., S. J. Counce, and D. F. Paulson, 1975 Synaptonemal complex complement of man in spreads of spermatocytes, with details of the sex chromosome pair. Science 187: 363–5.

Page, S. L., and R. S. Hawley, 2004 The genetics and molecular biology of the synaptonemal complex. Annu Rev Cell Dev Bi 20: 525–558.

Paigen, K., and P. Petkov, 2010 Mammalian recombination hot spots: properties, control and evolution. Nature Reviews Genetics 11: 221 233.

Paigen, K., J. P. Szatkiewicz, K. Sawyer, N. Leahy, E. D. Parvanov et al., 2008 The recombinational anatomy of a mouse chromosome. (M. Lichten, Ed.). PLoS Genetics 4: e1000119.

Pan, Z., Q. Yang, N. Ye, L. Wang, J. Li et al., 2012 Complex relationship between meiotic recombination frequency and autosomal synaptonemal complex length per cell in normal human males. Am J Med Genet A 158A: 581–587.

Peichel, C. L., S. R. McCann, J. A. Ross, A. F. S. Naftaly, J. R. Urton et al., 2020 Assembly of the threespine stickleback Y chromosome reveals convergent signatures of sex chromosome evolution. Genome Biol 21: 177.

Peichel, C. L., J. A. Ross, C. K. Matson, M. Dickson, J. Grimwood et al., 2004 The Master Sex-Determination Locus in Threespine Sticklebacks Is on a Nascent Y Chromosome. Curr Biol 14:1416-1424.

Rexroad, C. E., Y. Palti, S. A. Gahr, and R. L. Vallejo, 2008 A second generation genetic map for rainbow trout *(Oncorhynchus mykiss)*. BMC Genet 9: 74.

Ritz, K. R., M. A. F. Noor, and N. D. Singh, 2017 Variation in Recombination Rate: Adaptive or Not? Trends Genet 33: 364–374.

Ross, J. A., and C. L. Peichel, 2008 Molecular Cytogenetic Evidence of Rearrangements on the Y Chromosome of the Threespine Stickleback Fish. Genetics 179: 2173–2182.

Ruiz-Herrera, A., M. Vozdova, J. Fernández, H. Sebestova, L. Capilla et al., 2017 Recombination correlates with synaptonemal complex length and chromatin loop size in bovids—insights into mammalian meiotic chromosomal organization. Chromosoma 126: 615–631.

Sardell, J. M., C. Cheng, A. J. Dagilis, A. Ishikawa, J. Kitano et al., 2018 Sex Differences in Recombination in Sticklebacks. G3 8: 1971–1983.

Sardell, J. M., and M. Kirkpatrick, 2019 Sex Differences in the Recombination Landscape. Am Nat 195: 361–379.

Schneider, C. A., W. S. Rasband, and K. W. Eliceiri, 2012 NIH Image to ImageJ: 25 years of image analysis. Nat Methods 9: 671–675.

Segura, J., L. Ferretti, S. Ramos-Onsins, L. Capilla, M. Farré et al., 2013 Evolution of recombination in eutherian mammals: insights into mechanisms that affect recombination rates and crossover interference. Proc Royal Soc B Biological Sci 280: 20131945.

Shanfelter, A. F., S. L. Archambeault, and M. A. White, 2019 Divergent fine-scale recombination landscapes between a freshwater and marine population of threespine stickleback fish. Genome Biol Evol 11: 1573–1585.

Singer, A., H. Perlman, Y. Yan, C. Walker, G. Corley-Smith et al., 2002 Sex-specific recombination rates in zebrafish (Danio rerio). Genetics 160: 649–57.

Smeds, L., C. F. Mugal, A. Qvarnström, and H. Ellegren, 2016 High-Resolution Mapping of Crossover and Non-crossover Recombination Events by Whole-Genome Re-sequencing of an Avian Pedigree. PLoS Genetics 12: e1006044.

Smukowski, C. S., and M. A. F. Noor, 2011 Recombination rate variation in closely related species. Heredity 107: 496–508.

Solari, A. J., 1974 The Behavior of the XY Pair in Mammals. Int Rev Cytol 38: 273–317.

Solari, A. J., 1970 The spatial relationship of the X and Y chromosomes during meiotic prophase in mouse spermatocytes. Chromosoma 29: 217–236.

Soriano, P., E. A. Keitges, D. F. Schorderet, K. Harbers, S. M. Gartler et al., 1987 High rate of recombination and double crossovers in the mouse pseudoautosomal region during male meiosis. Proc National Acad Sci 84: 7218 7220.

Sun, F., M. Oliver-Bonet, T. Liehr, H. Starke, E. Ko et al., 2004 Human male recombination maps for individual chromosomes. Am J Hum Genet 74: 521–31.

Tease, C., and M. A. Hultén, 2004 Inter-sex variation in synaptonemal complex lengths largely determine the different recombination rates in male and female germ cells. Cytogenet Genome Res 107: 208 215.

Villena, F. P.-M. de, and C. Sapienza, 2001 Recombination is proportional to the number of chromosome arms in mammals. Mamm Genome 12: 318–322.

Wang, R. J., B. L. Dumont, P. Jing, and B. A. Payseur, 2019 A first genetic portrait of synaptonemal complex variation. Plos Genet 15: e1008337.

White, M. A., J. Kitano, and C. L. Peichel, 2015 Purifying Selection Maintains Dosage-Sensitive Genes during Degeneration of the Threespine Stickleback Y Chromosome. Mol Biol Evol 32: 1981–1995.

Wong, A. K., A. L. Ruhe, B. L. Dumont, K. R. Robertson, G. Guerrero et al., 2010 A comprehensive linkage map of the dog genome. Genetics 184: 595 605.

