## Supplementary material for "Sexual dimorphism of synaptonemal complex length among populations of threespine stickleback fish": Figure S1

**Figure S1.** The effect of hypotonic solution on synaptonemal complex length in males. Total length of SMC3 label along the synaptonemal complex was measured at late pachytene in males treated with either sodium citrate or sucrose as the hypertonic solution. The total number of measured fish is indicated below the plots. Whiskers denote 1.5x the interquartile range.

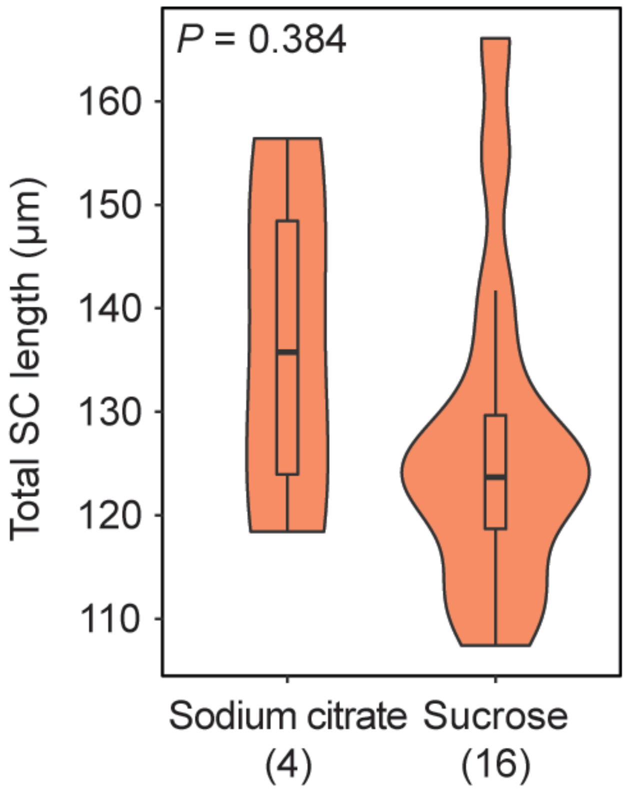
